# A systems biology-based identification and *in vivo* functional screening of Alzheimer’s disease risk genes reveals modulators of memory function

**DOI:** 10.1101/2022.08.19.504537

**Authors:** Adam D. Hudgins, Shiyi Zhou, Rachel N. Arey, Coleen T. Murphy, Yousin Suh

## Abstract

Genome-wide association studies (GWAS) have uncovered over 40 genomic loci associated with risk for late-onset Alzheimer’s Disease (LOAD), but identification of the underlying causal genes remains challenging. While the role of glial biology in the mediation of LOAD genetic risk has been increasingly recognized, recent studies of induced pluripotent stem cell (iPSC)-derived neurons from LOAD patients have demonstrated the existence of neuronal cell-intrinsic functional defects, absent interactions with other brain cell types or exposure to neurotoxic insults. Here, we searched for genetic contributions to neuronal dysfunction in LOAD pathobiology, using an integrative systems approach that incorporated multi-evidence-based gene-mapping and network analysis-based prioritization. We found widespread dysfunction in neuronal gene co-expression networks in the LOAD brain and identified synaptic and endolysosomal function as being specifically impacted by LOAD-associated genetic variation. A systematic perturbation screening of candidate risk genes in *C. elegans* revealed that neuronal knockdown of the LOAD risk gene orthologs *vha-10* (*ATP6V1G2*), *cmd-1* (*CALM3*), *amph-1* (*BIN1*), *ephx-1* (*NGEF*), and *pho-5* (*ACP2*) significantly alters short/intermediate-term memory function, the cognitive domain affected earliest during LOAD progression. These results highlight the impact of LOAD risk genes on evolutionarily conserved memory function, as mediated through neuronal endosomal dysfunction, and identify new targets for further mechanistic interrogation.

## Introduction

Alzheimer’s Disease (AD), the most common cause of dementia, is an age-related neurodegenerative disorder that affects millions worldwide^1^. Although our understanding of the molecular mechanisms underpinning the progression of AD has increased steadily over the past several decades^2,3^, the precise etiology of the disease remains elusive, and no preventative or curative treatments currently exist^3,4^. In recent years, large consortia-based genome-wide association studies (GWAS) have identified over 40 genomic loci associated with risk for sporadic late-onset AD (LOAD)^5–8^, the predominant form (>90% of cases) of the disease. However, the majority of risk variants reside in non-coding regions of the genome and are enriched in cell type-specific transcriptional regulatory elements such as enhancers, suggesting that they contribute to genetic risk by altering gene expression regulatory networks^9,10^. Yet, we still have a limited ability to predict how non-coding variants affect cell- and tissue-specific gene regulatory interactions that alter transcriptional outputs, confounding efforts to identify LOAD-causal variants and target genes^10–12^, a critical step to fully realize the promise of GWAS for clinical applications.

Thus far, the genes and loci implicated in LOAD genetic risk have nominated multiple pathways for disease relevance, including endosomal trafficking, cholesterol regulation, mitochondrial function, and inflammation and immunity^13^, most of which are active in multiple cell types in the brain. Due to the discovery of LOAD-associated coding variation in genes such as *TREM2*^14–16^, *PLCG2*^14,17^, and *ABI3*^14^, which are predominantly expressed in microglia, functional studies in both cell and animal models have increasingly been focused on the role of microglial biology with regard to genetic risk for LOAD and the pathways listed above^18–22^. However, in contrast to previous work which had highlighted the importance of microglia-expressed genes to transcriptional network dysregulation in the LOAD brain^23^, a recent co-expression network study of brain RNA-seq data from a large-scale LOAD cohort found neuron-specific co-expression modules to be the most profoundly affected by disease state^24^. Additionally, *in vitro* studies of neuronal cultures derived from LOAD patient iPSCs have demonstrated several cell-intrinsic defects in neuronal function, including hyperexcitability and altered synapse formation dynamics, absent interactions with other cell types or exposure to external neurotoxic insults^25,26^. The precise mechanisms through which common genetic variation contributes to neuronal cellular dysfunction and genetic risk for LOAD are understudied and remain largely unknown.

Here, we searched for genetic contributions to neuronal dysfunction in LOAD pathobiology by taking a systems biology approach. We analyzed summary statistics data from a recent LOAD GWAS meta-analysis^7^ in the context of large-scale brain omics data, utilizing 1) multi-evidence-based gene-mapping; 2) transcriptome-wide correlation with clinical and neuropathological traits and network analysis-based prioritization; and 3) *in vivo* functional screening to identify high-confidence neuronal genes and pathways contributing to LOAD pathophysiology. We found that many candidate LOAD risk genes that are dysregulated in the LOAD brain and more strongly correlated with clinically-assessed cognitive function and dementia severity than with post-mortem assessment of neuropathological burden are central members of network modules involved in critical neuronal functions.

As modeling cognitive dysfunction *in vitro* presents considerable challenges, we chose to screen our candidate genes for LOAD-relevant effects *in vivo*, through the use of *C. elegans* associative memory assays, a well-established experimental paradigm of cognitive function assessment with evolutionarily conserved molecular underpinnings^27–29^. *C. elegans* shares similarities with mammals in age-related physiological changes, including learning and memory decline^28^. Like mammals, memory loss is one of the earliest features of neuronal aging in *C. elegans*^28,30^. Furthermore, conserved molecular machinery is required in *C. elegans* to learn and remember^27,28,31^. A systematic perturbation screening of candidate risk genes in *C. elegans* revealed that neuronal knockdown of the LOAD risk gene orthologs *vha-10* (*ATP6V1G2*), *cmd-1* (*CALM3*), *amph-1* (*BIN1*), *ephx-1* (*NGEF*), and *pho-5* (*ACP2*) significantly altered short/intermediate-term memory function, the cognitive domain affected earliest during LOAD progression, highlighting these genes for further *in vitro* and *in vivo* evaluation as potential therapeutic targets.

## Results

### Integrative multi-omics analysis for target gene identification and functional screening *in vivo*

To identify high-confidence target genes that underlie LOAD genetic risk and contribute to neuronal dysfunction in LOAD pathobiology, we analyzed LOAD GWAS summary statistics^7^ in the context of large-scale brain omics data, as outlined in **Fig. 1**. Our analysis framework incorporates data from large-scale brain expression quantitative trait loci (eQTL) studies (PsychENCODE^32^, CommonMind Consortium (CMC)^33^, BRAINEAC^34^, BrainSeq^35^, ROSMAP^36^, and GTEx^37^), chromatin interaction data from various Hi-C analyses of brain and neural tissue (PsychENCODE - Dorsolateral Prefrontal Cortex (DLPFC)^32^, Giusti-Rodríguez et al.- Adult and Fetal Cortex^38^, and Schmitt et al. - DLPFC, Hippocampus, and Neural Progenitor Cell^39^), and RNA-seq data from a cohort of 364 brains from the Mount Sinai Brain Bank (MSBB)^40^, a recently generated resource made publicly available as part of the Accelerating Medicines Partnership-Alzheimer’s Disease Consortium (AMP-AD). A key strength of our approach is the use of the *C*. *elegans* short/intermediate-term associative memory assay as an organismal level readout of the relevance of our prioritized candidate genes to neuronal circuit integrity and function.

**Figure 1.**
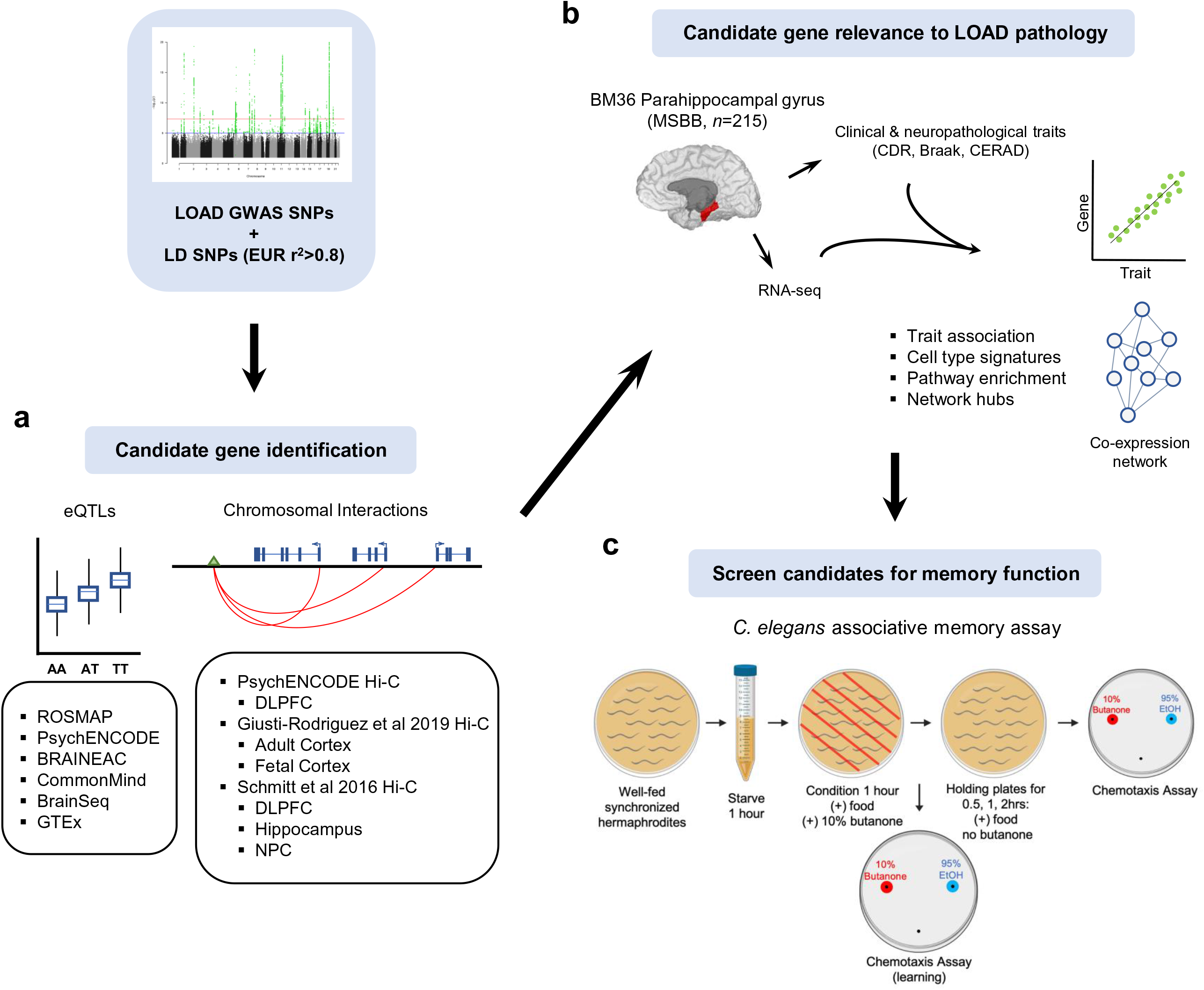
Integrative systems biology approach for LOAD risk gene identification and functional screening. (**a**) Candidate risk genes are identified from LOAD GWAS summary statistics, using functional genomics data from large-scale brain eQTL and chromatin interaction studies. (**b**) Relevance of candidate risk genes to LOAD biology is assessed by correlation of expression patterns with clinical and neuropathological traits, and connectivity within co-expression networks built from LOAD cohort brain RNA-seq data. (**c**) Prioritized candidate risk genes are screened for *in vivo* effects on memory function through the use of associative memory assays in *C. elegans*.

### LOAD GWAS variants are enriched in neuronal open chromatin regions

Previous studies have shown that LOAD SNP heritability is specifically enriched in transcriptional regulatory elements active in microglia^41–44^, findings which have contributed to the recent focus on microglial biology in LOAD functional genomics studies. While clearly important to genetic risk for LOAD, this microglial enrichment does not explain the dysfunctional phenotypes observed in LOAD patient iPSC-derived neurons^25,26^. To test for the presence of LOAD GWAS signal^7^ in neuronal transcriptional regulatory elements, we used single-cell open chromatin profiles generated from the human brain by Assay for Transposase-Accessible Chromatin using sequencing (scATAC-seq)^43^, and a statistical enrichment methodology employed by Wang and colleagues^45^ (see Methods). We found an enrichment of LOAD GWAS signal in the open chromatin of several neuronal cell types, over a wide range of statistical significance, from genome-wide significant (GWS) p-values (*P*<5×10^-8^) to a sub-GWS p-value of *P*=1×10^-4^, although the overall neuronal enrichment observed was much weaker than that seen in microglia (**Fig. 2a**).

**Figure 2.**
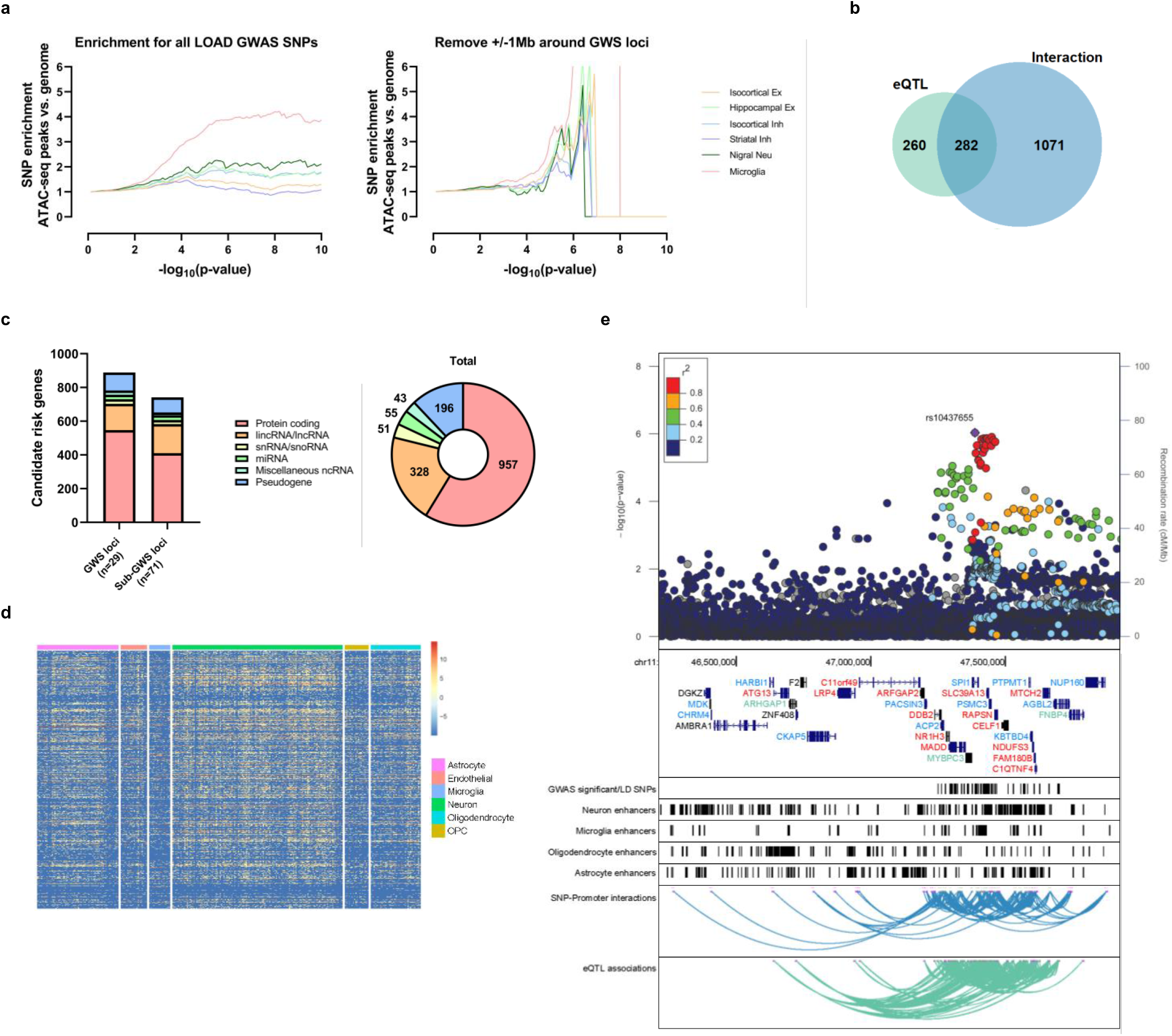
Data from eQTL and chromatin interaction studies implicates potential causal genes in LOAD GWAS loci. (**a**) Enrichment signal for sub-threshold LOAD GWAS SNPs in neuronal open chromatin becomes evident following the removal of GWS loci and nearby SNPs (+/- 1 Mb), becoming similar in magnitude to that of microglia. Each point on the curves represents the difference in fold of the proportion of SNPs with a p-value below the cutoff in the ATAC-seq peaks versus all SNPs present in the GWAS summary statistics. (**b**) Numbers of candidate risk genes unique to, and shared by, the two gene-mapping methods. (**c**) Distribution of candidate risk genes by gene type and significance threshold. (**d**) Heatmap of cell type-specific expression patterns of candidate risk genes in the human brain. Color scale represents relative expression across cell types (red = higher, blue = lower). (**e**) Example LOAD GWAS locus (*CELF1*/*SPI1*), highlighting challenges in the identification of causal genes. Top to bottom – Manhattan plot of -log10(p-value) association statistics from Jansen et al., with the top SNP rs10437655 highlighted in purple and remaining variants colored according to LD (*r*^2^) with the lead SNP; Genome browser track showing all coding genes present in the locus. Gene names colored in green or blue are candidate risk genes nominated by QTL evidence or SNP-promoter interaction evidence, respectively. Gene names colored in red are candidate risk genes nominated by both kinds of evidence; Track showing the location of significant GWAS SNPs (*P*<1×10^-5^), and SNPs in LD (*r*^2^>0.6); Tracks indicating the positions of enhancer elements identified in different human brain cell types; Track illustrating the significant chromatin interactions between LOAD GWAS SNPs and gene promoters in the locus; Track illustrating the significant eQTL associations between LOAD GWAS SNPs and genes in the locus.

Since any enrichment of signal in the sub-GWS range could potentially be explained by linkage disequilibrium (LD) with above-threshold LOAD GWAS variants^9^, we performed an additional enrichment analysis after removing all variants within 1 Mb of the GWS loci. Surprisingly, the enrichment of sub-GWS signal in neuronal open chromatin regions was significantly strengthened, a result observed for all neuronal subtypes in the scATAC-seq dataset, including both excitatory and inhibitory neurons (**Fig. 2a**). In comparison, the same analysis using scATAC-seq data from the human lung^46^ did not show enrichment in the open chromatin of lung cell types (**Supplemental Fig. 1a**). This result indicates that sub-threshold LOAD GWAS loci likely harbor causal non-coding risk variants in transcriptional regulatory elements active in neurons, which may lead to the dysregulation of causal risk genes underlying the dysfunction observed in LOAD patient iPSC-derived neurons. Thus, for our integrative approach outlined in **Fig. 1** we chose to include loci which reached a suggestive significance threshold of *P*<1×10^-5^, in addition to GWS loci, as this approach has been successfully used for post-GWAS gene mapping^45,47–49^.

### eQTL and chromatin interaction data indicate potential causal genes in LOAD GWAS loci

Increasing evidence suggests that the gene nearest to the most significant variant in a GWAS loci is often not the causal gene^50–53^. To identify and prioritize candidate causal LOAD risk genes, we incorporated functional genomics data with summary statistics from the recent LOAD GWAS meta-analysis conducted by Jansen and colleagues^7^ using the web-based platform Functional Mapping and Annotation (FUMA)^54^ (see Methods). We selected genes which were nominated by eQTL^32–37^ or chromatin interaction data^32,38,39^ (**Fig. 1**) and disregarded genes that were only implicated through positional mapping. These datasets expand upon those used for gene-mapping in the original Jansen et al. study^7^, incorporating two more eQTL studies (PsychENCODE^32^ and BrainSeq^35^), and two additional Hi-C studies (PsychENCODE^32^ and Giusti-Rodríguez et al.- Adult and Fetal cortex^38^). In addition, we included those genes that contained protein-coding variants in LD (*r^2^* > 0.8) with the tag variant at each LOAD GWAS locus. This strategy nominated 1,630 coding and non-coding genes, in 29 GWS and 71 sub-GWS loci, as candidate causal LOAD risk genes (**Fig 2c, Supplemental Table 1**). The majority of mapped genes were protein-coding, with lncRNAs and pseudogenes making up the next two largest categories, in roughly equal proportions in both GWS and sub-GWS loci (**Fig. 2c**).

More candidate risk genes were mapped by variant-promoter chromatin interactions (n=1,353) than by eQTL evidence (n=542). In total, 282 genes (17%) were supported by both eQTL and chromatin interaction evidence (**Fig. 2b**). The PsychENCODE and the Adult and Fetal cortex Hi-C data provided most of the chromatin interaction-implicated genes (**Supplemental Fig. 2a, Supplemental Table 2**), and the majority of cis-eQTL-associated genes came from the PsychENCODE, GTEx, and CMC datasets (**Supplemental Fig. 2b, Supplemental Table 3**). Using human single-cell transcriptome data from the temporal cortex^32,55,56^, we examined the expression patterns of the protein-coding candidates (n=957) and found ubiquitous expression across cell types for many genes, but higher levels of expression for most of the risk genes in neurons and astrocytes, in comparison to that of microglia, oligodendrocytes, oligodendrocyte precursors, and endothelial cells (**Fig. 2d**).

Of particular interest is the *CELF1/SPI1* locus (11p11.2), which did not reach GWS in the Jansen et al. study^7^ but has been found as GWS in several previous LOAD GWAS^5,6,8^ (**Fig. 2e**). Previous work has implicated *SPI1* as the causal gene in the locus^57^ based on the data from microglia. However, whether or not *SPI1* is the only causal gene in the locus, and microglia are the only causal cell type, remains unclear. Overlaying the locus with recently generated brain cell type-specific epigenomic annotation data^42^, we found that the region of LOAD association is rich with regulatory elements that are active in several cell types, including dense clusters of neuronal enhancers (**Fig. 2e**). Our integrated analysis including eQTL and chromatin interaction data implicates almost all (n=29) of the protein-coding genes in the *CELF1/SPI1* locus as candidate causal LOAD risk genes (**Fig. 2e**), highlighting the challenges in identifying the true causal genes and relevant cell types underlying GWAS associations and the need for further prioritization and functional screening.

### Prioritized LOAD risk genes are co-expression network hubs dysregulated in the LOAD brain

To determine the potential relevance of our candidate risk genes to the transcriptional alterations occurring in the LOAD brain, we performed weighted gene co-expression network analysis (WGCNA)^58^ on RNA-seq data from a cohort of 364 brains from the Mount Sinai Brain Bank (MSBB)^40^. Samples from this cohort span a wide spectrum of LOAD-related neuropathological and cognitive disease severities; and RNA-seq data exists for 4 brain regions: Brodmann area 10 (BM10) frontal pole, BM22 superior temporal gyrus, BM36 parahippocampal gyrus, and BM44 inferior frontal gyrus. We focused our analyses on data from the BM36 region (n=215) because a prior transcriptomic study of this cohort found that BM36, out of 19 sampled regions, was the most highly affected by AD^59^. Using WGCNA we identified 32 distinct co-expression modules, 10 of which were enriched for cell type-specific gene expression signatures: oligodendrocyte (M1, M22); neuronal (M2, M10, M25, M32); astrocyte (M23); endothelial (M14, M18); microglia (M28); astrocyte/endothelial (M20); and microglia/endothelial (M21) (**Fig. 3a**).

**Figure 3.**
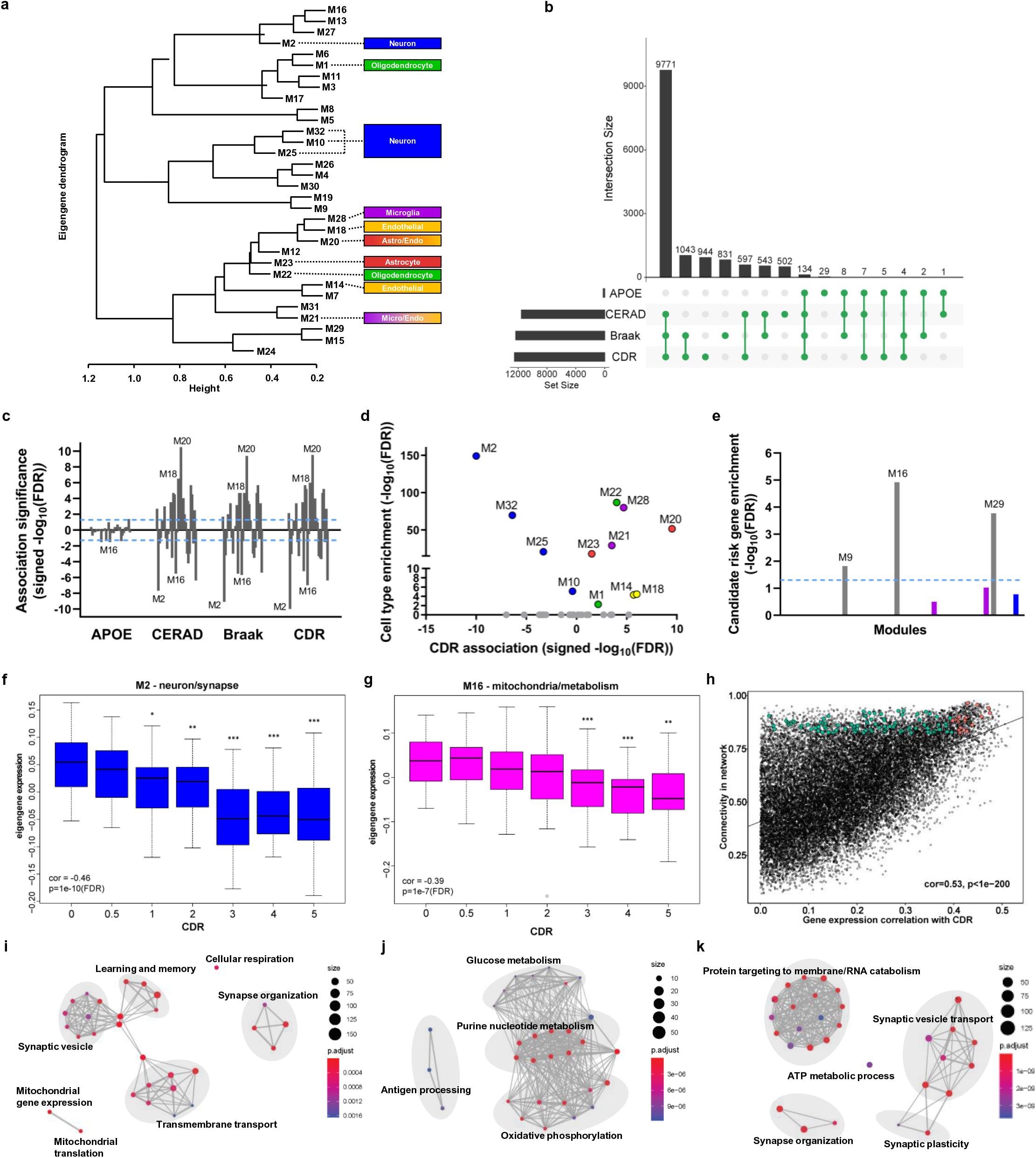
Co-expression network analysis identifies candidate LOAD risk genes as dysregulated neuronal subnetwork hubs in the LOAD brain. (**a**) Co-expression network analysis of RNA-seq data from the parahippocampal gyrus identifies 32 distinct co-expression modules. Modules enriched for cell type-specific gene expression signatures are indicated. (**b**) UpSet plot of the intersections between gene sets found to be significantly associated with the listed traits. (**c**) Association significance of correlations between module eigengenes and traits. Significance of the top four trait-associated modules (M2, M20, M16, M18) is indicated. Bars extending past the dotted line represent FDR < 0.05. (**d**) Scatter plot of module eigengene association with CDR vs. enrichment of cell type gene expression signature. Blue dots = neuronal modules, green dots = oligodendrocyte modules, red dots = astrocyte modules, purple dots = microglia modules, yellow dots = endothelial modules. (**e**) Significance of enrichment of LOAD candidate risk genes within each module. Grey bars = modules with no cell type enrichment, blue bars = neuronal modules, purple bars = microglia modules. Bars extending above the line represent FDR < 0.05. (**f**-**g**) Expression of the module eigengene decreases significantly with increased dementia severity for both the neuronal/synapse module M2 (**f**) and the mitochondrial/metabolism module M16 (**g**). Pearson’s correlation and FDR-corrected *P*-value are indicated. Differences in the expression of the module eigengene at each CDR score with respect to cognitive baseline (CDR=0) was also assessed by t test. (**h**) Gene expression correlation with CDR is significantly correlated with network connectivity as measured by kME. Pearson’s correlation and FDR-corrected *P*-value are indicated. Core network candidate risk genes, according to max kME, are shown in teal. The top 20 high-priority risk gene candidates, as determined by correlation with CDR and network centrality, are highlighted in orange. (**i**-**k**) Significantly enriched (FDR < 0.05) Gene Ontology biological process terms are shown as network maps, with edges connecting overlapping gene sets, for the neuronal/synapse module M2 (**i**), the mitochondrial/metabolism module M16 (**j**), and the core network genes (**k**). Node size indicates the number of genes overlapping with the term and node color indicates magnitude of adjusted *P-*value significance. \**P* < 0.05, \*\**P* < 0.01, \*\*\**P* < 0.001.

We utilized the spectrum of neuropathology and cognitive function present across the dataset to identify significant associations between gene expression and disease severity. We assessed the correlations between both the expression of individual genes, and the expression of module eigengenes, and neuropathological category (CERAD), neurofibrillary tangle burden (Braak), and clinically-assessed cognitive function (CDR), as well as *APOE* genotype. On the level of individual genes, 14,421 genes were associated with at least one trait (FDR<0.05, **Supplemental Table 4**), indicating large-scale rewiring of transcriptional activity in the LOAD brain, with a large overlap seen between genes that were significantly correlated with CDR score, CERAD neuropathological category, and Braak staging score (**Fig. 3b**). In contrast, *APOE* genotype was significantly associated with relatively few genes (**Fig. 3b**). With regard to module-trait correlations, we found that 26 of the 32 modules were significantly associated with at least one trait (**Fig. 3c**). The top four modules with the strongest associations were: the neuronal module M2, which was most strongly negatively associated with CDR score; the astrocyte/endothelial module M20, which was most strongly positively associated with CERAD category; module M16, which was not enriched for any cell type-specific signature but was the second-most negatively correlated with CDR score; and the endothelial module M18, which was the second-most positively associated with CERAD category (**Fig. 3c**). Modules which were enriched for cell type-specific signatures showed a clear demarcation with respect to their correlation with CERAD category, and Braak and CDR scores, with all neuronal modules having a negative association, while astrocyte, microglia, oligodendrocyte, and endothelial modules were all positively associated (**Fig. 3d, Supplemental Table 5**). We then assessed whether any of the modules were enriched for our candidate LOAD risk genes, considering only protein-coding genes which had a significant correlation with a trait (**Fig. 3e**). The module most enriched for our candidate risk genes was M16, one of the top four trait-associated modules, along with two other modules, which were also not enriched for any cell type-specific expression signature: module M19, which was negatively correlated with CDR; and module M29, which was positively correlated with Braak score (**Fig. 3e**).

Previous findings from network-based analysis of LOAD transcriptomic data have highlighted the disease-relevance of modules representing aspects of microglia^23^ and oligodendrocyte^60^ biology. Most recently, a co-expression network analysis of the same RNA-seq dataset we analyzed here, utilizing a different methodology, determined that neuronal modules were the most significantly affected by LOAD pathobiology^24^. A key difference and advantage of our study is the use of genetic association as the fundamental basis of our prioritization schema, upon which we leverage network approaches to derive new insights from LOAD brain transcriptome data. It has been recognized that genetically-supported drug targets have a much greater chance of success in clinical trials^61^. By using genetics as a foundation, we increase confidence in our prioritized risk genes while also increasing the probability of successful therapeutic development. Indeed, our analysis confirmed that the significant disease-relevant neuronal modules contain well-supported AD risk genes, including the familial AD gene *APP* and the APP processing pathway member *SORL1*^62,63^ in module M2, which was the most strongly associated with dementia status (CDR) (**Fig. 3c**). The overall expression of M2 member genes, as captured by the module eigengene, exhibited marked downregulation during the progression from normal cognitive function to advanced AD dementia (**Fig. 3f**). Correspondingly, gene ontology analysis of M2 member genes revealed an enrichment for many biological processes that are profoundly affected by AD, including synaptic signaling, learning and memory, synaptic structure and organization, transmembrane transporter activity, and regulation of mitochondrial transcription and translation (**Fig. 3i**). Another module of significance was M16, a top trait-associated module with the strongest enrichment for our candidate risk genes. In comparison to M2, M16 member genes didn’t display significant downregulation until more advanced levels of cognitive decline (**Fig. 3g**) and showed an enrichment for biological process terms encompassing many aspects of mitochondrial function, including oxidative phosphorylation, and glucose and purine nucleotide metabolism (**Fig. 3j**).

### Identification of high-priority candidate causal risk genes for functional screening *in vivo*

The network analysis identified important relationships between genes and modules involved in synapse and mitochondrial biology, critical components of healthy neuronal function, and LOAD. Since our candidate LOAD risk genes were enriched in these important neuronal function co-expression modules and were more closely correlated with dementia status than with neuropathological burden, we chose to functionally screen our candidates for effects on memory, in a non-amyloidosis model, in neurons *in vivo*. The *C*. *elegans* short/intermediate-term associative memory (S/ITAM) assay was chosen as the ideal experimental paradigm due to the highly evolutionarily conserved molecular biology which underpins memory function from worms to mammals^27–29^, as well as the practicality and efficiency the model affords, allowing for the testing of large numbers of candidates.

For an un-biased, systematic analysis, we included 4 categories of candidates (**Table 1**). 1) **High-priority candidate causal risk genes.** Since candidate risk genes with higher centrality in the network are more likely to have disease-relevant effects if perturbed^64–66^, we focused on candidates which occupied centrally-connected nodes within the overall co-expression network. We identified “core” genes as those positioned in the top 10% of the network, as determined by the eigengene-based connectivity measure kME (see Methods). Interestingly, overall connectivity in the co-expression network displayed a strong correlation with gene-trait association (**Fig. 3h**) so that genes with the highest absolute correlation with CDR score were more likely to have high network centrality. Furthermore, core genes were enriched for biological process terms involved in neuronal functions, including synaptic plasticity, synaptic vesicle transport, and synapse organization, as well as ATP metabolic processes, protein targeting to the membrane, and RNA catabolism (**Fig. 3k**). We ranked these core network candidate risk genes by absolute correlation with dementia status and prioritized the top 20 as high-priority targets for functional validation (**Supplemental Table 6**). Notably, eighteen of these top 20 candidates were either not the genes usually nominated from their respective loci or were genes that came from sub-GWS loci^6,7^. The two exceptions were the well-replicated LOAD GWAS gene *PTK2B*, and the familial AD gene *APP* (**Supplemental Table 6**), which resides in a locus that reached the suggestive association threshold (*P*<1×10^-5^)^7^. Out of the 20 high-priority candidates, 16 were members of the neuronal signature modules M2 and M32. The remaining four candidates were members of M16, the module with the strongest enrichment for candidate LOAD risk genes (**Fig. 3e**), and one of the top four trait-associated modules. 2) ***CELF1*/*SPI1* locus candidates**. Since the expression of 18 of the 29 eQTL- and chromatin interaction-implicated genes in the *CELF1*/*SPI1* locus (**Fig. 2e**) were significantly correlated with CDR score (**Table 1, Supplemental Table 4**), we selected multiple members (n=5) of this locus to screen for potential memory effects. 3) **Well-studied LOAD GWAS genes.** We selected two of the best studied GWS LOAD GWAS genes, *BIN1* and *PICALM*, based on recent fine-mapping analyses using cell type-specific approaches^42,43^ and their known role in synaptic function^67–69^, as strong candidates. 4) **Candidates unsupported by prioritization schema**. Since any effects on memory function we observed in our screen could conceivably occur due to perturbation of important neuronal genes that coincidentally exist in LOAD GWAS loci but have no actual relevance to LOAD genetic risk, we included genes from LOAD GWAS loci that were not prioritized by our analysis (*RAPSN* and *GRIN3B* (not present in MSBB RNA-seq data); *GNB2* and *TRPM7* (not correlated with CDR score); and *CHL1*, *GDE1*, and *RORA* (previously identified sub-GWS LOAD GWAS loci^5^ which did not meet significance criteria^7^)), to act as surrogate negative controls. To identify the appropriate targets for our 33 prioritized candidate LOAD risk genes (**Table 1, Supplemental Table 6**), we used the web-based comparative genomics tool OrthoList 2^70^ to identify the closest *C*. *elegans* orthologs for our perturbation screen. Keeping only those genes with orthology predictions supported by more than one database, we found 27 well-supported orthologs for 24 of our candidate risk genes (**Table 1**, **Supplemental Table 7**). As a final layer of prioritization, we selected orthologs which had been shown to be expressed in *C*. *elegans* neurons, as determined by our previous work characterizing the *C. elegans* neuronal transcriptome^71^ (**Table 1**, **Supplemental Table 7**), leading to a total of 27 worm orthologs of 24 LOAD GWAS candidate risk genes in 17 loci for *in vivo* functional screening.

### Neuron-specific knockdown of LOAD risk gene orthologs alters memory function in *C*. *elegans*

Since the expression of all our candidate genes were negatively correlated with LOAD severity, with the exception of *BIN1* and *PICALM* (**Table 1**), we knocked-down the candidate genes using RNAi to mimic the directional impact of association. To generate neuronal-specific knockdown of candidate genes, we used a neuronal RNAi-sensitive strain (LC108) of *C. elegans*, which can otherwise be refractory to RNA interference. We knocked-down each of the LOAD candidate risk gene orthologs from egg stage and tested for effects on short/intermediate-term memory (at 1 hour and 2 hours post-conditioning) at day 1 (young adulthood). Knockdown of most of the candidate genes had no effect on naive chemotaxis (**Supplemental Fig. 3a-g**), suggesting that they did not alter normal neuronal development or function, with the exceptions of *F54A3.2* (*CKAP5*; decreased naive chemotaxis), and *aps-2* (*AP2S1*; increased naive chemotaxis), which unfortunately prevented robust assessment of any potential memory effects for these two high-priority genes. In addition, *rpt-5* (*PSMC3*), *unc-11* (*PICALM*), and *gpb-1* (*GNB2*) could not be assayed for memory effects due to motor deficits resulting from knockdown. Finally, two of the high-priority candidates, *kin-32* (*PTK2B*) and *cisd-1* (*CISD1*), could not be tested for memory effects due to a lack of available RNAi clones in the Ahringer and Vidal libraries. However, their presence in our list of top candidates gave us further confidence in our prioritization schema because of previous findings from functional studies of these genes. In mice, *PTK2B*, a well-known LOAD GWAS gene, has been shown to have important roles in hippocampal-dependent memory, synaptic plasticity, and dendritic spine structure^72^, and deficiency of *CISD1*, a gene involved in mitochondrial function, has been shown to elicit Parkinsonian phenotypes^73^.

Among the high-priority candidates that could be tested in the memory assays (**Table 1**), knockdown of *vha-10* (*ATP6V1G2*), *cmd-1* (*CALM3*), and *ephx-1* (*NGEF*) caused significant impacts on memory function (**Fig. 4a, 4b, 4d**), while knockdown of *jnk-1* (*MAPK9*), *apl-1* (*APP*), *Y62E10A.2* (*POP7*), *misc-1* (*SLC25A11*), and the other *CKAP5* ortholog, *zyg-9*, had no effect (**Fig. 4a-c**). Knockdown of our top candidate *vha-10* (*ATP6V1G2*) resulted in a robust memory deficit at 1hr post-conditioning (**Fig. 4a**). *ATP6V1G2* encodes a neuronal-specific subunit of the vacuolar-type ATPase (V-ATPase), a proton translocating pump that plays critical roles in the acidification of endosomal compartments including lysosomes^74^ and the loading and release of synaptic vesicles^75^. Similarly, knockdown of *cmd-1* (*CALM3*), encoding the calcium-binding protein calmodulin, resulted in a significant memory deficit at 1hr and 2hr post-conditioning (**Fig. 4b**). In the brain calmodulin has diverse functions, including the regulation of synaptic signaling, endocytosis, cholesterol metabolism, and ion channel function^76^. Interestingly, knockdown of the high-priority risk candidate *ephx-1* (*NGEF*) and the well-known LOAD GWAS risk gene *amph-1* (*BIN1*) had no effect on short/intermediate-term memory function, but instead resulted in increased memory retention at 2hr post-conditioning **(Fig. 4d**). These results indicate that neuronal loss of expression of these genes impacts processes of active forgetting that are mediated through RAC1/CDC42^77,78^. Indeed, *NGEF* encodes a neuronal guanine nucleotide exchange factor (GEF) that regulates the activity of GTPases such as RAC1, RHOA, and CDC42^79^, and recent work has implicated *BIN1* in RAC1-mediated synaptic remodeling^80^.

**Figure 4.**
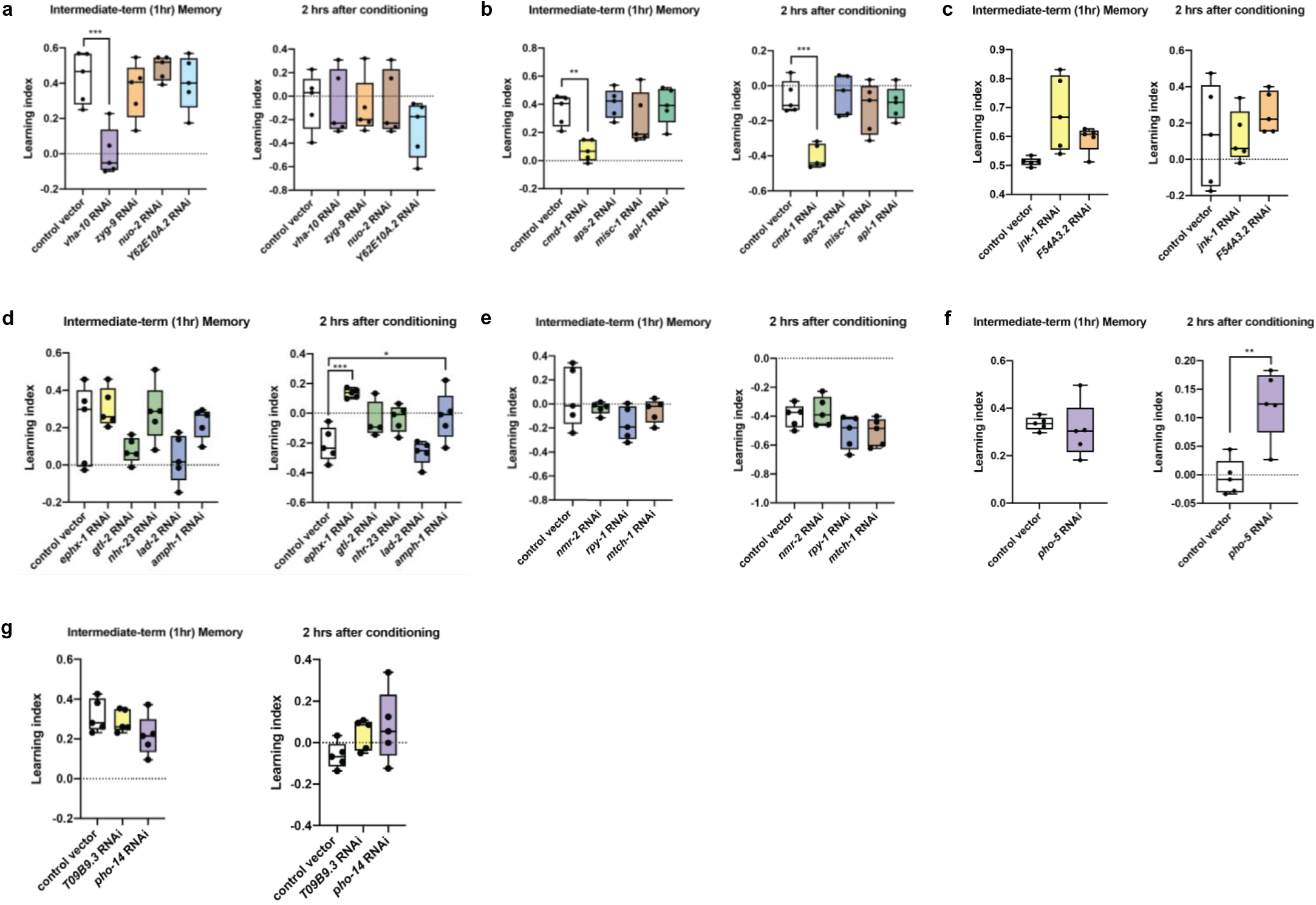
Neuronal knockdown of LOAD risk gene orthologs alters memory function in *C. elegans*. (**a**-**g**) 1 hour and 2 hour post-conditioning learning indices of worms treated with whole-life RNAi for LOAD candidate risk gene orthologs. Grouping of the tested orthologs was random and does not represent candidate prioritization. n ≥ 4 (n: technical replicates). Statistical significance determined by One-way ANOVA, with Dunnett’s post hoc test. \**P* < 0.05, \*\**P* < 0.01, \*\*\**P* < 0.001, \*\*\*\**P* < 0.0001.

Among the 5 gene orthologs from the gene dense *CELF1*/*SPI1* locus (**Table 1**) that did not meet the criteria for inclusion in the list of high-priority candidates, *mtch-1* (*MTCH2*), *nuo-2* (*NDUFS3*), *rpt-5* (*PSMC3*), and two orthologs of *ACP2*, *pho-5* and *pho-14,* knockdown of *mtch-1*, *nuo-2*, and *pho-14* showed no significant memory effects (**Fig. 4a, 4e, 4g**). However, knockdown of *pho-5*, the closest ortholog of the lysosomal acid phosphatase gene *ACP2,* resulted in a memory retention effect at 2hr post-conditioning, similar to what we observed for *ephx-1* and *amph-1* (**Fig. 4f**). This result suggests that, like *NGEF* and *BIN1*, *ACP2* is also involved in the process of active forgetting, possibly through the local turnover of synaptic proteins during dendritic spine remodeling, a process recently found to involve neuronal-activity dependent lysosome trafficking^81^.

In total, out of 24 LOAD GWAS candidate risk genes (27 worm orthologs) in 17 loci tested, we identified 5 genes in 5 loci as *in vivo* modulators of memory function (**Table 1**). Taken together, the results of our systematic perturbation screen indicate that LOAD genetic risk impacts neuronal function, particularly with respect to memory, through two primary avenues – the synapse (*ATP6V1G2*, *CALM3, BIN1*, *NGEF*), and the lysosome (*ACP2*, *ATP6V1G2, CALM3*). The common point of interaction between these two fundamental components of neuronal biology is the endosomal trafficking system. Pathway and gene set analyses have previously found a significant enrichment of LOAD-associated genetic variation in genes involved in endolysosomal function^82^. However, the involvement of the endolysosomal system in LOAD pathobiology has typically been conceptualized in the context of amyloid and tau biology^83–85^. Our findings indicate that genetic contributions to neuronal dysfunction in LOAD pathobiology can affect the endolysosomal system through mechanisms which do not involve amyloid and tau, but instead directly impact the evolutionarily conserved pathways of learning and memory.

## Discussion

Studies of the genetic underpinnings of LOAD continue to uncover new genomic loci of interest but identifying the responsible genes and translating genetic discoveries into druggable targets remains a major challenge for the field. In this study we searched for the genetic contributions which underlie neuronal dysfunction in LOAD pathobiology, using an integrative systems approach that incorporated multi-evidence-based gene-mapping and network analysis-based prioritization, with the *C*. *elegans* short/intermediate-term associative memory assay as an organismal level readout of the impact of our prioritized candidate risk genes on neuronal circuit integrity and function.

We compiled and employed a large array of functional genomics data to identify candidate risk genes from LOAD GWAS loci. Examined in the transcriptional context of the LOAD brain, we found significant associations between many candidate risk genes and phenotypic measures of cognitive dysfunction and LOAD neuropathology. Network analysis identified several neuronal co-expression modules that were the most significantly associated with LOAD-associated cognitive dysfunction. We prioritized candidate risk genes by using genetic association and functional genomics evidence, focusing on core genes in the neuronal co-expression modules. A limitation of functional genomics-enabled post-GWAS gene mapping is the possibility of false positive gene nominations due to such factors as non-causal overlap between QTL and GWAS associations and non-disease relevant promiscuous chromatin interactions between GWAS variants and gene promoter regions. This limitation persists regardless of the quality and comprehensiveness of the tools and datasets used. Because of this fact, functional follow-up is critical to gaining confidence in a set of GWAS-implicated genes. Thus, we conducted functional studies of the prioritized candidate neuronal risk genes, as well as low-priority and non-prioritized genes, for effects on *in vivo* memory function in *C*. *elegans*.

Testing 27 worm orthologs out of 24 LOAD GWAS candidate risk genes in 17 loci, this study is to our knowledge the first comprehensive functional screen of its kind. The most notable finding of this study is the identification of 5 LOAD causal risk genes, *ATP6V1G2*, *CALM3*, *BIN1*, *NGEF*, and *ACP2*, in 5 loci, as *in vivo* modulators of evolutionarily conserved memory function. *ATP6V1G2* encodes a neuronal-specific subunit of the large V-ATPase complex. Our analysis prioritized *ATP6V1G2* as our top candidate risk gene, both due to its membership within the core network of genes, as well as being the candidate most significantly associated with cognitive function. *ATP6V1G2* has not been previously nominated as a LOAD risk gene, most likely due to the fact that it resides in the 6p21.32 major histocompatibility (MHC) locus, a region well-known for having an extremely complex LD structure that makes the identification of causal variants and genes in the locus particularly difficult. Multiple members of the V-ATPase complex are associated with neurological disorders and neurodegenerative conditions arising due to defective lysosomal acidification^86^. Additionally, V-ATPase function has also been shown to be important for the maintenance of neural stem cell renewal capacity^87^, and the age-related loss of this capacity is also implicated in impaired cognitive function^88^. In support of our findings, a network-based study found *ATP6V1A* (3q13.2), another member subunit of V-ATPase, to be one of the top drivers of neuronal function that is dysregulated in the LOAD brain^24^. Furthermore, testing for LOAD relevance in a *Drosophila* model of Aβ pathology, the authors of the study^24^ found that *Vha68-1* (*ATP6V1A*) deficiency negatively affected neuronal activity and exacerbated Aβ-mediated neuronal toxicity. These findings complement our observation that *vha-10* (*ATP6V1G2*) deficiency causes deficits in short/intermediate-term memory function in *C*. *elegans*, and further highlights evolutionarily conserved V-ATPase function as an attractive target for LOAD therapeutic development.

*CALM3* is one of the three identical isoforms of the calmodulin gene that is encoded in the human genome. A calcium-binding factor, calmodulin is ubiquitously expressed, and has central roles in a wide variety of processes critical to cellular health and function. Calmodulin function has been tied to LOAD pathobiology for some time, leading some to postulate a “calmodulin hypothesis” for AD pathogenesis^89^, as an extension of the already-established “calcium hypothesis”^90^. With respect to *CALM3* in particular, it has been difficult to definitively tie alterations in *CALM3* expression in the LOAD brain to genetic risk because *CALM3* resides within the greater *APOE* locus. Due to the powerful LOAD association of *APOE*, along with the strong LD relationships in this locus, identification of additional signals beyond the well-studied *APOE* coding variants^91–93^ has been challenging. We identified and prioritized *CALM3* by our analyses for *in vivo* testing. In contrast to mammals, the *C. elegans* genome contains only one ortholog of calmodulin, *cmd-1*. We found that neuron-specific knockdown of this critical gene *cmd-1* (*CALM3*) resulted in a significant memory deficit at 1hr post-conditioning without causing significant motor or chemotaxis defects. This interesting phenotype likely involves differential regulation of calmodulin-dependent kinase II (CaMKII) activity, given its well-known roles in learning, memory, and forgetting^94,95^.The *BIN1* (Bridging Integrator 1) locus has the second strongest LOAD association behind *APOE*. Recent variant fine-mapping studies have indicated that transcriptional regulatory elements specific to microglia might be the mediators of LOAD genetic risk in the region, resulting in altered microglial *BIN1* expression^42,43^. However, different cell types in the brain express different isoforms of *BIN1*, and while global transcription of *BIN1* is increased in the LOAD brain, the transcription of neuron- and astrocyte-specific isoforms are downregulated and are associated with tau pathology^96^. Additionally, recent work has shown that neuron-specific conditional knockout of *BIN1* results in reduced synapse density, decreased presynaptic vesicle release, and learning and memory deficits in mice^67^. Interestingly, we found that neuronal-specific knockdown of the *C*. *elegans* ortholog *amph-1* (*BIN1*) resulted in a decreased ability to “forget” an associated memory, in line with its role in RAC1-mediated synaptic remodeling^80^, an important component in the process of active forgetting^77,78^. These results suggest complex roles for neuronal BIN1 function that may have isoform-dependent phenotypes upon perturbation.

*NGEF*, or ephexin-1, is a neuronal guanine nucleotide exchange factor (GEF) for GTPases such as RAC1, RHOA, and CDC42. Besides central functions in axon guidance^79^ it also has major roles in dendritic spine morphogenesis, post-synaptic organization, and pre-synaptic vesicle release through its interactions with Eph receptors like EphA4^97,98^. We found that, similar to *amph-1* (*BIN1*), neuronal knockdown of *ephx-1* (*NGEF*) resulted in a persistence of associative memory. In the LOAD GWAS locus that includes *NGEF*, *INPP5D* is the gene usually nominated as causal. However, a recent fine-mapping study identified neuron-specific chromatin interactions between LOAD risk variants and the *NGEF* promoter^42^, nominating *NGEF* as one of the top candidate neuronal causal genes. These results indicate that there might be multiple, cell type-specific, causal genes in this locus, in contrast to the prevailing view that LOAD genetic risk is conferred by dysregulation of *INPP5D* primarily in microglia^99^.

The *CELF1/SPI1* LOAD GWAS locus (11p11.2), which we screened extensively for functional effects on memory, is a gene dense locus that did not reach GWS in the Jansen et al. 2019 GWAS/X meta-analysis^7^, but has been found as GWS in several previous studies^5,6,8^. *SPI1* has been found to be a likely causal gene with regard to the relevance of this locus for microglial function in AD^57^, but LD relationships in this locus are complex and other lines of evidence^42,43,100^ as well as the results presented here indicate that this locus harbors additional causal genes, including *ACP2*. *ACP2* encodes lysosomal acid phosphatase 2, a phosphatase present in the lysosomal membrane which assists in the maturation of lysosomal enzymes and helps maintain the optimal pH for proper lysosomal function^101^. In humans *ACP2* is broadly expressed in all tissues, with particularly strong expression in pyramidal neurons of the cortex and cerebellar Purkinje cells^102^. *ACP2* deficiency in mice causes lysosomal storage defects, seizures, skin, cerebellum, and vertebral malformations, and ataxia^103,104^. Intriguingly, a recent LOAD whole exome sequencing (WES) study identified a rare missense variant in *ACP2* (D353E) to be enriched in controls compared to LOAD patients^105^, suggesting a protective role of *ACP2* in LOAD. Correspondingly, we found that neuron-specific knockdown of *pho-5* (*ACP2*) results in extended associative memory in *C*. *elegans*, even up to 3 hours post-conditioning, an interesting result which agrees directionally with the finding from the WES analysis. While complete loss of ACP2 function results in severe neurological phenotypes^103,104^, these results suggest that reduced ACP2 function could be protective with respect to LOAD-associated cognitive impairment.

In addition to endolysosomal biology, mitochondrial function was also enriched in our top LOAD-associated neuronal modules, and both have been implicated in the etiology of other neurodegenerative diseases, including Parkinson’s disease (PD)^106^. Interestingly, several PD risk genes are members of the modules we highlight in this study, including *GBA* and *PINK1* (mitochondrial module M16), and *SNCA* and *PRKN* (neuronal module M2). Additionally, gene ontology analysis of the LOAD-downregulated mitochondrial function module M16 found a significant enrichment for genes involved in antigen presentation (**Fig. 3k**), and recent studies have drawn links between mitochondrial antigen presentation and immune responses in PD^107^, suggesting potentially common mechanisms of pathogenesis between the two diseases, centered on mitochondrial biology. Notably, a previous co-expression network study found two modules which were conserved between normal aging and LOAD, one representing mitochondrial processes, and the other representing synaptic function, and identified *ATP6V1G2* as a top hub gene common to both LOAD and aging^108^. Since modules and genes that we identified through our work have also been found to be relevant to the normal aging process, this suggests that perhaps LOAD genetic risk factors which affect neuronal function are the earliest contributors to disease pathophysiology, as aging is the greatest risk factor for neurodegenerative disease, including LOAD.

In summary, our integrative analysis and *in vivo* screening revealed genetic contributions to neuronal dysfunction in LOAD pathobiology and identified evolutionarily conserved key neuronal genes and pathways involved in this process. When combined with the growing publicly available human genomic data, simple model organism systems, such as the *C. elegans* behavioral paradigm used here, have great potential to advance the functional genetic understanding of the complex etiology of LOAD.

## Acknowledgments

We thank the Murphy and Suh lab members for their input. This work was supported by NIH grants AG057433, AG061521, AG055501, AG057706, AG057909, and AG017242 (Y.S.) The work was also supported by NIH grant AG057341 to C.T.M. and Y.S..

## Author Contributions

Y.S. and C.T.M conceptualized the study. A.D.H., S.Z., and R.N.A. performed experiments and analyzed data. A.D.H., S.Z., R.N.A., C.T.M. and Y.S. wrote the manuscript.

## Methods

### Data sources

Alzheimer’s disease GWAS summary statistics from the Jansen et al. meta-analysis^7^ were retrieved from https://ctg.cncr.nl/software/summary_statistics. The MSBB LOAD RNA-seq data are available through the AD Knowledge Portal (https://adknowledgeportal.synapse.org) under the synapse ID# syn3159438. Processed single-cell expression data from human brain^32,55,56^ was downloaded from the PsychENCODE Integrative analysis web portal (http://resource.psychencode.org), as listed under the descriptor “Processed single-cell expression data merged from all three sources”. Human brain cell type-specific enhancer tracks from Nott et al.^42^ are available through the UCSC Genome Brower (https://genome.ucsc.edu/s/nottalexi/glass_lab_BrainCellTypes_hg19).

### LOAD patient cohort

The Mount Sinai Brain Bank (MSBB) LOAD cohort consists of 364 postmortem control and LOAD patient brains, each accompanied by robust clinical and neuropathological phenotype metadata, with various sample subsets used for the generation of genome-, transcriptome- and proteome-scale molecular datasets, as has been described in detail previously^40^. For our analyses we utilized bulk RNA-seq data that had been generated from the Brodmann area 36 parahippocampal gyrus region of a subset of the greater cohort (n=215). Each individual had full neuropathological assessments according to the Consortium to Establish a Registry for Alzheimer’s Disease (CERAD) protocol^109^, a Braak staging score for neurofibrillary neuropathology burden^110^, and a Clinical Dementia Rating (CDR) scale score^111^ based on premortem dementia and cognitive function assessment. *APOE* genotype was available for a subset of the individuals (n=135).

### Candidate gene mapping

To map LOAD GWAS loci to genes we used the web-based tool Functional Mapping and Annotation (FUMA, v1.3.6a)^54^. Using the summary statistics from the Jansen et al. meta-analysis, a sub-genome-wide significance threshold of *P*<1×10^-5^ was used to identify all independent (*r*^2^<0.1, EUR population, 1000 Genomes) loci. Within each identified locus, all SNPs that met the significance threshold were used for mapping, as well as SNPs in strong linkage disequilibrium (*r*^2^>0.6, EUR population, 1000 Genomes) with the index variant of each locus. Gene mapping was conducted using two strategies: 1) Selecting genes with significant cis-eQTL associations (FDR < 0.05) with the LOAD GWAS SNPs (i.e., expression of the gene is associated with allelic variation at the SNP). Six large-scale brain eQTL studies were utilized for this purpose – PsychENCODE^32^, CommonMind Consortium^33^, BRAINEAC^34^, BrainSeq^35^, ROSMAP^36^, and GTEx v8 (Brain and Nerve tissue only)^37^; and 2) Selecting genes by identifying significant chromatin interactions (FDR ≤ 1e-6) between gene promoter regions (250 bp up- and 500 bp downstream of the transcription start site) and the LOAD GWAS SNPs, as identified by Hi-C data. Data from three Hi-C studies of brain and neural tissue were utilized – DLPFC from PsychENCODE^32^, Adult and Fetal cortex data from the study of Giusti-Rodríguez et al.^38^, and DLPFC, Hippocampus, and Neural Progenitor Cell data from the study of Schmitt et al.^39^.

### Cell type expression specificity of candidate risk genes

We investigated the cell type-specific expression of our candidate risk genes by examining their expression patterns in published human single-cell transcriptome data from the temporal cortex^32,55,56^. Preprocessed single-cell expression data^32,55,56^ was downloaded as described above, and only the adult, broad cell class data was retained (Astrocyte, Endothelial, Microglia, Neuron, Oligodendrocyte, OPC). Expression data was scaled and log-normalized and displayed as a heatmap by using the R package ‘pheatmap’.

### Statistics

Descriptions of all statistical tests performed are included in the figure legends or the respective Methods sections, where relevant.

### Co-expression network analysis

The R package ‘WGCNA’^58^ was used to construct a co-expression network from the MSBB LOAD brain RNA-seq data^40^. For the creation of the network we utilized the publicly available preprocessed expression matrix (see Data sources) which had already been normalized and adjusted for sex, race, age, RNA integrity, post-mortem interval, and batch effect. A weighted co-expression network was built using the preprocessed expression values and the blockwiseModules WGCNA function with the following parameters: soft-thresholding power = 8, TOMType = “signed”, deepSplit = 2, minimum module size of 15, merge cut height of 0.25, signed hybrid network with pamRespectsDendro = FALSE. This resulted in the identification of 32 modules of co-expressed genes, from which we calculated module eigengenes (MEs). Correlations between clinical and neuropathological traits and individual gene expression or MEs were computed as Pearson’s correlations and were corrected for multiple testing according to the FDR (Benjamini-Hochberg) method. Significance was determined using an adjusted *P*-value cutoff of 0.05.

### Enrichment analysis

Enrichment of LOAD GWAS signal^7^ in open chromatin regions of human brain cell types was calculated according to the methodology of Wang et al.^45^, using called peaks from scATAC-seq of the human brain^43^. At each p-value significance cut-off, using a sliding -log(p-value) threshold from 0 to 10 in steps of 0.1, the proportion of SNPs in ATAC-seq peaks with p-values more significant that the cut-off, the foreground, was calculated against the proportion of SNPs present in the summary statistics (∼13 m). Co-expression modules were tested for significant overlap with the cell type-specific expression signatures of five major brain cell types (neurons, microglia, astrocytes, oligodendrocytes, endothelial) identified through human brain single-cell RNA-sequencing data^55^, and reported previously^59^. Enrichment statistics were calculated by one tailed Fisher’s exact test and corrected for multiple comparisons by the FDR (Benjamini-Hochberg) method. Significance was determined using an adjusted *P*-value cutoff of 0.05. Functional enrichment of biological pathways within the co-expression modules was assessed by over-representation test, using the R package ‘clusterProfiler’^112^, considering all genes present in the MSBB RNA-seq dataset as the set of background genes. The Gene Ontology (Biological Process) gene sets used for the enrichment analysis came from the Molecular Signatures Database (MSigDB) v7.0^113,114^. Multiple testing correction was performed according to the FDR (Benjamini-Hochberg) method and significance was determined using an adjusted *P*-value cutoff of 0.05. Significantly enriched terms were visualized as a network map, with edges connecting overlapping gene sets, using the emapplot function of the R package ‘enrichplot’^115^, with layout = “kk”.

### Identification of core network genes

To identify genes with high trait-relevance that also reside in centrally-located positions within the co-expression network, we took the approach used by Chateigner et al.^116^. By utilizing the module membership measure kME (the correlation between the expression of a gene and the module eigengene), it can be appreciated that the genes with the highest kME in a given module are also the most correlated to the traits that are most closely associated with the module eigengene. This relationship demonstrates the utility of employing kME as a centrality score when prioritizing genes with both relevance to the trait of interest and high network connectivity. Using kME to define the topological positions of all the genes in the co-expression network, the max kME was identified for every gene (i.e., the score with respect to the module to which the gene was assigned), and “core” network genes were then defined as the top 10% of genes with the highest global absolute scores.

### Worm cultivation

All strains were maintained at 20°C on plates made from high growth medium (HGM: 3 g/L NaCl, 20 g/L Bacto-peptone, 30 g/L Bacto-agar in distilled water, 4 mL/L cholesterol (5 mg/mL in ethanol), 1 mL/L 1M CaCl2, 1 mL/L 1M MgSO4, and 25 mL/L 1M potassium phosphate buffer (pH 6.0) added to molten agar after autoclaving (Brenner, 1974) with OP50 *E.coli* as the food source. For RNAi treatment, the standard HGM was supplemented with 1 mL/L 1M IPTG (isopropyl b-d-1-thiogalactopyranoside) and 1 mL/L 100 mg/mL carbenicillin, and plates were seeded with HT115 *E. coli* for ad libitum feeding. Worms were synchronized by collecting eggs from hermaphrodites via exposure to an alkaline-bleach solution (80 mL water, 5 mL 5N KOH, 15 mL sodium hypochlorite); collected eggs were repeatedly washed in M9 buffer (6 g/L Na2HPO4, 3 g/L KH2PO4, 5 g/L NaCl and 1 mL/L 1M MgSO4 in distilled water; Brenner, 1974).

### Strains

LC108 (*vIs69 [pCFJ90(Pmyo-2::mCherry + Punc-119::sid-1*)])

### Short/intermediate-term associative memory training

Worms were tested for short/intermediate-term memory as previously described (Kauffman et al., 2010). Briefly, synchronized day 1 adult hermaphrodites were washed from HGM plates with M9 buffer for 3 times. Then the animals were starved for 1 hr in M9 buffer. For training, worms were transferred to 10 cm NGM conditioning plates seeded with OP50 E. coli bacteria and with 18 ul 10% 2-butanone (Acros Organics) in ethanol on the lid for 1 hr. After conditioning, the trained worms were tested for chemotaxis towards 10% butanone vs. an ethanol control either immediately (0 hr) or after being transferred to 10 cm NGM plates with fresh OP50 for specified intervals before testing (30 min-2 hr). Chemotaxis indices (CI) were calculated as follow: (#wormsButanone – #wormsEthanol)/(Total #worms). Learning indices (LI) are: LI_trained=_CI_trained_-CI_naive_.

---

**Supplemental Figure 1. Lack of enrichment of LOAD GWAS SNPs in open chromatin of lung cell types**

(**a**) Enrichment signal for LOAD GWAS SNPs in open chromatin of lung cell types, both with the inclusion of GWS loci and following the removal of GWS loci and nearby SNPs (+/- 1 Mb). Each point on the curves represents the difference in fold of the proportion of SNPs with a p-value below the cutoff in the ATAC-seq peaks versus all SNPs present in the GWAS summary statistics.

**Supplemental Figure 2. More candidate risk genes were mapped by variant-promoter chromatin interactions than by eQTL evidence**

(**a**) UpSet plot of the intersections between gene sets nominated by the chromatin interaction data from various Hi-C analyses of brain and neural tissue. (**b**) UpSet plot of the interactions between gene sets nominated by the large-scale brain expression quantitative trait loci (eQTL) studies.

**Supplemental Figure 3. Naive chemotaxis is mostly unaffected after neuronal knockdown of LOAD risk gene orthologs in *C. elegans***

(**a**-**g**) Naive chemotaxis indices of worms treated with whole-life RNAi for LOAD candidate risk gene orthologs. Grouping of the tested orthologs was random and does not represent candidate prioritization. n ≥ 4 (n: technical replicates). Statistical significance determined by One-way ANOVA, with Dunnett’s post hoc test. \**P* < 0.05, \*\**P* < 0.01, \*\*\**P* < 0.001, \*\*\*\**P* < 0.0001.

