## Supplementary figures and images for "A systems biology-based identification and *in vivo* functional screening of Alzheimer’s disease risk genes reveals modulators of memory function"

### Supplemental Figures

#### Supplemental Fig. 1

**a**

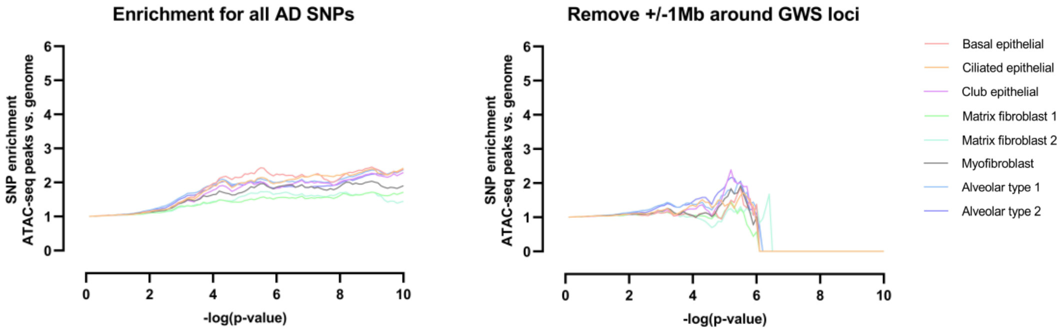

#### Supplemental Fig. 2

a b

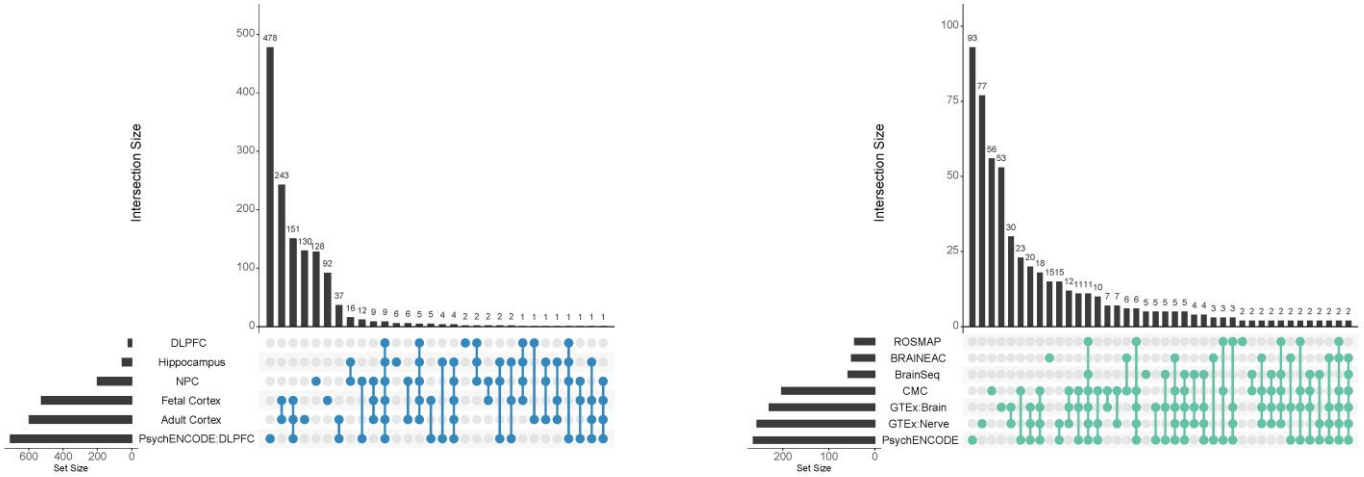

#### Supplemental Fig. 3

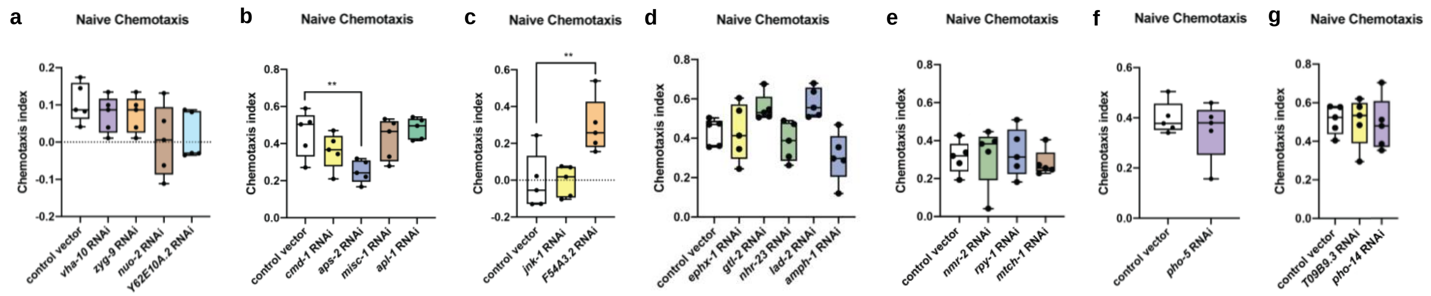
